# Epistasis facilitates the long-term antigenic evolution of the influenza B virus hemagglutinin

**DOI:** 10.64898/2026.07.22.739994

**Authors:** Lara S. U. Schwab, Ruopeng Xie, Ellie Reilly, Malet Aban, Shu Hu, Natalie Spirason, Yi-Mo Deng, Randy Suryadinata, Monica Galiano, Matthew J. Gartner, Kanta Subbarao, Karen Laurie, Steve Rockman, Stephen J. Kent, Adam K. Wheatley, Ian G. Barr, Vijaykrishna Dhanasekaran, Marios Koutsakos

## Abstract

The antigenic drift of viral glycoproteins must be balanced by purifying selection pressure to maintain functionality. Understanding these evolutionary processes is key to predicting and combating viral evolution but is primarily based on influenza A(H3N2), which may limit generalisability. By characterising the influenza B virus haemagglutinin (HA) over 8 decades of circulation in humans, we found continuous genetic diversification, punctuated with antigenic changes that did not follow a linear path in antigenic space. Antigenic change is primarily underpinned by re-occurring mutations and deletions at positions 136, 150, 162-165, 197 and 203. These residues form complex epistatic networks that modulate the antigenic impact of mutation recycling. They also generate permissive backbones on which immune escape can emerge with limited replicative fitness cost. Our study identifies critical similarities and differences with A(H3N2) evolution and demonstrates the role of epistasis in balancing antigenic novelty with viral fitness. Our findings and genetic, antigenic and phenotypic datasets support the development of genotype-to-phenotype prediction tools, but such predictions need to capture the complex outcomes of epistasis.

## Introduction

Viral surface glycoproteins that mediate attachment and entry have critical roles in the life cycle and are thus the target of neutralising antibodies. Circulating viruses face opposing selective pressures: maintaining glycoprotein function for viral replication while accumulating mutations to escape population-level immunity(1–3). For viruses that continually circulate in human populations like influenza, these opposing selective pressures create a dynamic evolutionary landscape where viral fitness and antigenic innovation must be balanced(3, 4). Understanding how mutations in viral glycoproteins are facilitated and constrained is fundamental to predicting viral evolution and developing durable vaccines. Our understanding of these evolutionary processes primarily comes from studies of influenza A(H3N2)(5–9). A(H3N2) viruses evolve rapidly along a relatively linear trajectory in antigenic space, with a relatively small number of amino acid residues in the proximity of the receptor binding site contributing to antigenic change(6, 9). There is evidence that epistasis (whereby the effect a mutation depends on the genetic backbone on which it occurs) modulates the antigenic impact of mutations(10, 11) as well as imposing entrenchment towards specific evolutionary paths by modulating the fitness cost of mutations(12–15). While these findings have revealed how the antigenic evolution of the A(H3N2) hemagglutinin (HA, major surface glycoprotein of influenza viruses that mediates attachment and entry) proceeds under different selection pressures, the generalisability of these mechanisms remains unclear. Indeed, the evolutionary patterns and constrains of even relatively closely related viruses (like the different influenza A subtypes) can differ considerably(16, 17). Thus, while the study of A(H3N2) has established a foundational framework of viral evolution, it remains important to consider other viruses to establish generalisable rules of viral evolution.

In the more than 80 years since their discovery, human influenza B viruses (IBV) have undergone substantial genetic and antigenic diversification, including the emergence of two antigenically distinct and co-circulating lineages since around the 1980s, designated B/Victoria (after B/Victoria/2/87) and B/Yamagata (after B/Yamagata/16/88)(18). Both lineages co-circulated globally for over four decades, until the disappearance of B/Yamagata post-2020(19, 20). A(H3N2) and IBV exhibit marked differences in mutational tolerance, evolutionary rates and circulation dynamics(16, 21, 22), making IBV an ideal model system to further probe the long-term antigenic evolution of viral surface glycoproteins and test the generalisability of inferences from A(H3N2). IBV evolution has been less extensively studied than A(H3N2) or A(H1N1), despite its clinical and public health relevance(18, 23–26), accounting for approximately one-quarter (20-33%) of annual influenza cases(25). Based on our current understanding of IBV HA antigenicity, various broadly acting vaccine candidates have been designed(27), with the hope of mitigating the clinical burden of IBV or the, at least theoretical, possibility of eradicating this virus(28). However, for such vaccines to be broadly protective, or even universal, the different variants of IBV and the molecular basis of their emergence should be defined.

The antigenic structure of IBV HA, which comprises of 4 antigenic regions in the HA head (29), has been mapped primarily through escape mutant selection using monoclonal antibodies. These studies have identified 4 distinct antigenic sites (the 120 loop, 150 loop, the 160 loop and the 190 helix) comprising approximately 30 amino acid residues surrounding the receptor binding site (RBS). More recently, integration of historical sequence and antigenic data via phylogenetic analysis (NextFlu/Nextstrain) identified 33 specific amino acid substitutions across B/Yamagata and B/Victoria lineages that correlated with antigenic changes(30). This number of antigenic residues is much larger than those defined for A(H3N2)(6), however, the vast majority of these positions/sites have not been experimentally validated for their antigenic impact on the IBV HA. Indeed, only mutations at positions 148, 149, 150, and 203 have been experimentally shown (by site directed mutagenesis and reverse genetics) to alter the antigenic phenotype of the IBV HA, and this was specifically in the context of the B/Victoria and B/Yamagata lineage emergence(31). Thus, while sequence analysis suggests many positions may be important for antigenic change, the molecular determinants of antigenic cluster transitions and trajectories in IBV have remained experimentally unvalidated. In addition, such analyses have been restricted to the more recent IBV evolution and have not considered the entirety of IBV evolution over the last 8 decades. Overall, a greater understanding of the IBV antigenic evolution and its molecular basis could inform surveillance, bi-annual vaccine strain selection and next-generation vaccine design.

To address these gaps and to test the generalisability of inferences from A(H3N2) evolution, we phylogenetically and experimentally characterised the evolution of the IBV HA over 81 years of human circulation (1940–2021) and determined the molecular basis of antigenic drift during this period. We aimed to: (i) reconstruct the antigenic evolution of IBV HA over eight decades using serological data and computational antigenic mapping; (ii) identify amino acid residues that drive antigenic cluster transitions within each IBV lineage through reverse genetic analysis and hemagglutination inhibition (HAI) assays; (iii) validate antigenic effects using human sera to ensure biological relevance; (iv) assess the replicative fitness of antigenic variants to understand how viral fitness is maintained during antigenic evolution; and (v) define epistatic interactions that enable antigenic innovation while preserving viral function. Our genetic, antigenic, and phenotypic datasets span three IBV lineages and 12 antigenic clusters, providing an unprecedented foundation for understanding long-term influenza B haemagglutinin evolution and potentially informing new influenza vaccine development strategies.

## Results

### Evolution of the IBV HA from 1940-2021

To characterise the long-term evolution of IBV HA, we analysed 405 HA sequences from isolates between 1940 and 2021. These sequences predominantly originated from Asia (26.3%), Australasia (36.2%) or Europe (26.8%), with 52% of them derived from cell-propagated virus isolates (Figure S1a-b). Time scaled phylogenetic analysis confirmed a single pre-1970s lineage circulating until the early 1970s (previously referred to as Ancestral or Early(31, 32), when the first B/Victoria-lineage virus (B/Hong Kong/5/1972) was detected (Figure 1a)(31). Consistent with previous work(31, 33), two phylogenetically distinct IBV lineages circulated in the 1970s and early 1980s, one giving rise to modern-day B/Victoria viruses and the other giving rise to the B/Yamagata lineage.

**Figure 1.**
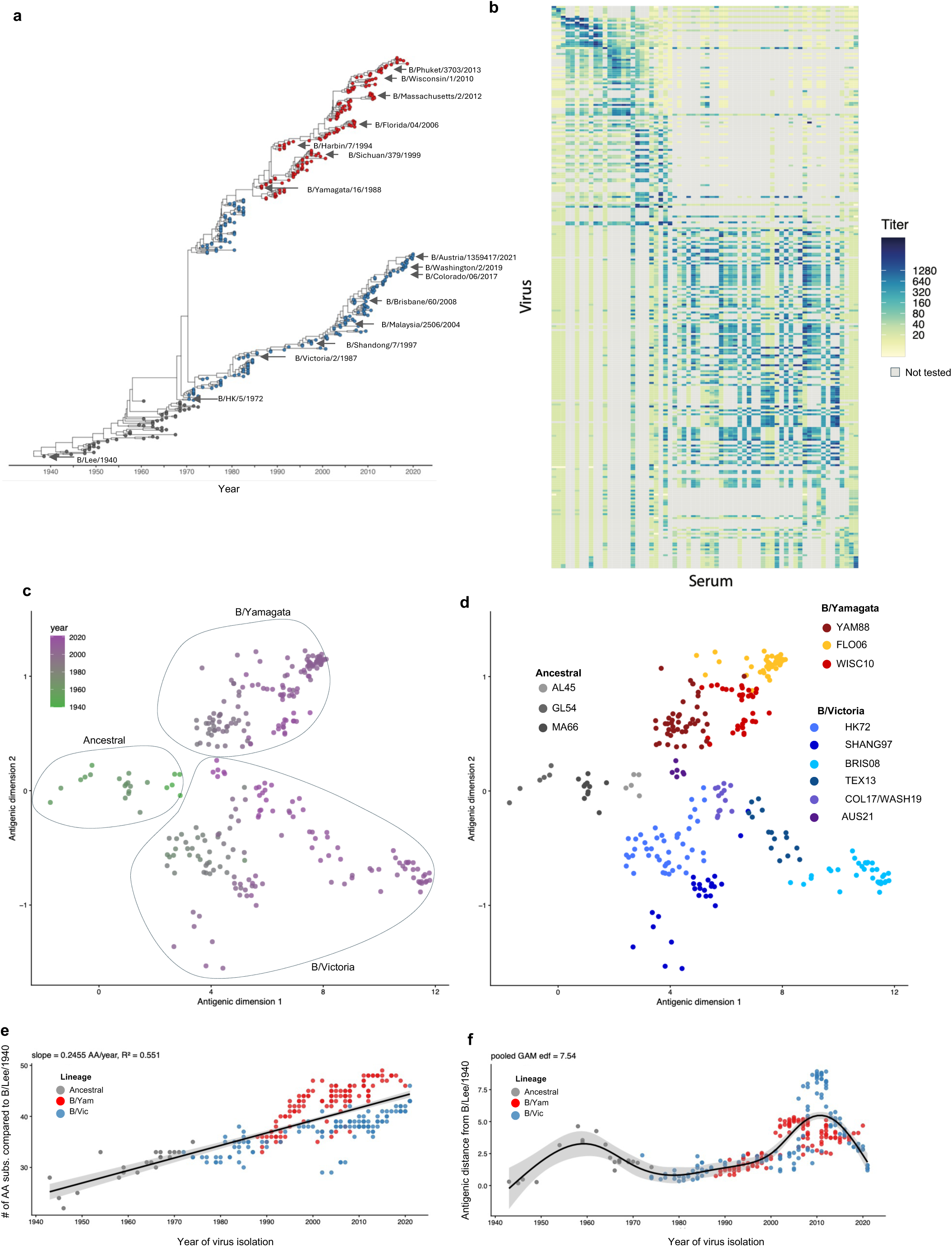
Genetic and antigenic evolution of the IBV HA between 1940-2021. (a) Maximum clade credibility (MCC) tree of IBV coloured by the antigenic lineage of the HA. Representative vaccine strains are labelled. (b) Heat map of HAI data used for antigenic cartography. Rows represent viruses and columns represent antisera. Colour intensity indicates HI titre, with grey denoting untested virus–antiserum combinations. (c) Antigenic evolution of IBV HA inferred using Bayesian multidimensional scaling (BMDS). Each circle represents the estimated antigenic location of an individual virus in two-dimensional antigenic space and is coloured by year of isolation. Internal nodes represent inferred ancestral antigenic states, and grey lines connect ancestral and descendant nodes according to the MCC tree, illustrating the evolutionary trajectory of antigenic change. (d) Same BMDS antigenic map as shown in c, coloured by antigenic cluster. (e) Number of amino acid substitutions of each IBV isolate compared to B/Lee/1940, plotted against the year of virus isolation. (f) Antigenic distance of each IBV isolate from B/Lee/1940 plotted against the year of virus isolation.

Selection pressure analyses revealed pervasive purifying selection across most codons, with episodic positive selection identified at 20 sites (Table S1). Several residues, including codon 197, were consistently supported by multiple methods across the full dataset, ancestral viruses, and both B/Victoria and B/Yamagata lineages, whereas other sites (including 150, 163, 165 and 203) exhibited method-or dataset-specific signals of selection. In addition, Bayesian Graphical Models (BGM) identified 85 coevolving pairs of sites with posterior probability greater than 0.5. Many of these coevolving pairs involve residues within or proximal to known antigenic regions and the receptor-binding site, as well as interactions between surface-exposed and structurally constrained residues, suggestive of epistatic and compensatory evolution.

### Antigenic evolution of the IBV HA from 1940-2021

To understand the antigenic evolution of the IBV HA, we constructed an antigenic map of the IBV HA using HAI titres of 287 IBV isolates and 66 ferret antisera from 1940-2021 (Figure 1b, S1c) (details in methods). Antigenic data were analysed jointly with HA sequence data using a Bayesian multidimensional scaling (BMDS) model, enabling simultaneous reconstruction of antigenic phenotypes and evolutionary relationships while quantifying antigenic and genetic distances among strains. The resulting antigenic map captured the antigenic diversification of the two lineages and indicated a non-linear path for B/Victoria viruses through antigenic space (Figure 1c). We identified 12 antigenic clusters, comprising 3 clusters within the Ancestral lineage, 6 within B/Victoria, and 3 within B/Yamagata. Antigenic evolution primarily proceeded along the first antigenic dimension, with discrete transitions between clusters (Figure 1d).

Despite the continuous accumulation of amino acid substitutions at an estimated rate of 0.246 amino acids per year (Figure 1e), antigenic distance did not increase linearly away from B/Lee/1940 (Figure 1f). Before 2000, both B/Victoria and B/Yamagata lineages exhibited relatively slow antigenic drift. Estimated rates were 0.087 units per year (95% CI 0.064 to 0.107) in the B/Victoria HK72 cluster and 0.130 units per year (95% CI 0.103 to 0.156) in the B/Yamagata YAM88 cluster. Between 2000 and 2005, antigenic evolution accelerated during the transition to the B/Victoria SHAN97 cluster, which was associated with acquisition of a B/Yamagata NA gene, and during the emergence of the B/Yamagata FLO06 cluster. Estimated rates during this period were 0.217 (95% CI 0.054 to 0.379) and 0.279 (95% CI 0.163 to 0.395) antigenic units per year, respectively. After 2005, B/Yamagata viruses transitioned to the WISC10 cluster and exhibited minimal antigenic change, with an estimated mean rate of −0.040 antigenic units per year, indicating near stability along the first antigenic dimension. In contrast, B/Victoria viruses showed a marked acceleration in antigenic evolution between 2005 and 2010, reaching 1.103 antigenic units per year (95% CI 0.696 to 1.510) within the BRIS08 cluster. From 2010 to 2021, antigenic drift in B/Victoria continued at a rate of −0.615 antigenic units per year (95% CI −0.665 to −0.565), reflecting a directional shift along the first antigenic dimension and spanning multiple clusters, including TEX13, COL17/WASH19, and AUS21. Notably, the most recent AUS21 cluster occupied an antigenic position close to Ancestral viruses, indicating convergence toward earlier antigenic states, consistent with previous report(31). Overall, we found punctuated antigenic drift, and an antigenic reversion of recent B/Victoria viruses.

### Antigenic impact of egg passage

Egg passage of influenza viruses can introduce HA mutations that may impact antigenicity. This includes the loss of glycosylation site motif (197N-198E-199T) on the IBV HA, although the antigenic impact of this glycosylation change is only partially understood(34, 35). Due to the historic nature of the viruses in our study going back 8 decades, unavoidably some viruses were isolated in eggs (Figure S1b). Of the egg-grown isolates in our HA sequences, 89.9% carried a mutation that disrupted the 197-199 glycosylation site (e.g. N197D, T198I). The remaining 10% of viruses carried the G141R mutation, which prevents the loss of glycosylation in eggs(34). To determine the antigenic impact of egg passage in our dataset, we firstly compared the HAI titres of antisera from ferrets inoculated with 3 matched cell or egg grown IBV against a panel of IBV isolates. HAI titres were strongly correlated, although for one of the viruses (B/Brisbane/60/2008), antisera raised against the egg isolate had greater HAI activity against certain antigens (Figure S2a). Similarly, when testing a panel of ferret antisera against matched cell or egg grown IBV antigens, HAI titres were strongly correlated, but the egg-grown B/Brisbane/60/2008 antigen could be recognised more strongly by some antisera. (Figure S2b). When comparing their positions in our antigenic map, matched cell or egg grown antigens (n=14) were on average only 0.47 antigenic units apart. Overall, the impact of egg adaptation appears specific to some IBV isolates but not others. We note that the recent appearance of N197D in B/Victoria viruses from ∼2020 onwards(30) indicates that this glycosylation change cannot be considered purely an egg adaptive mutation but one that also occurs in nature.

### Molecular basis of antigenic change

To understand the molecular basis of BHA antigenic evolution we compared the HA sequences of consecutive antigenic clusters and identified amino acid substitutions between successive clusters within each lineage. As the mutations that underpinned the emergence of the B/Victoria lineage from the Ancestral lineage and the emergence of B/Yamagata from B/Victoria were previously characterised by Rosu et al(31), we focused on the additional antigenic cluster transitions within each lineage. We chose representative viruses from each cluster for which ferret anti-sera were available and then used reverse genetics to introduce single or multiple point mutations in the HA. Viruses were tested in HAI assay using ferret antisera from the initial and subsequent cluster (Figure S3a). We considered a mutation to have an effect if the HAI activity of at least one of the ferret anti-sera changed by 2-fold or more.

Within the Ancestral lineage, inhibition of B/Allen/1945 by its homologous antiserum was reduced by mutations K136I/T, N150K and R203I/T but not N75K or V170L. These mutations only partially increased inhibition by the B/GL/1954 antiserum (Figure S3b). Inhibition of B/GL/1954 by its homologous or the related B/Maryland/1/1959 antiserum were reduced by I203T but not the T121A or K241Q mutations, but none of them increased inhibition by the B/Mass/66 antiserum (Figure S3c).

Within the B/Victoria lineage, the cluster transition between B/Malaysia/2506/2004 and B/Brisbane/60/2008 was underpinned by the N165K mutation, but not N75K or S172P (Figure S4a). The B/Colorado/06/2017 and B/Washington/02/2019 viruses formed one cluster in our antigenic map, but we assessed them separately as they (i) are phylogenetically distinct, (ii) involve different potential mutations and (iii) were considered antigenically distinct for vaccine strain selection(36). Inhibition of B/Brisbane/60/2008 by its homologous antiserum was reduced by the 2aa deletion (Δ162-163), although only by 2-fold, while the N129G mutation had no effect on its own (Figure S4b). Inhibition of B/Brisbane/60/2008 by its homologous antiserum was also reduced by the 3aa deletion (Δ162-164) and K136E, but not by G133R. All three mutations increased recognition by the B/Washington/02/2019 antiserum (Figure S4c).

The most recent B/Victoria cluster (B/Austria/1359417/2021) arose from B/Brisbane/60/2008 with Δ162-164 and K136E and an additional 8 mutations, including N150K, K203R and N197D (Figure S5a). As the B/Brisbane HA reverse genetics likely originated from an egg-passaged virus(37) and thus already contained N197D, we reverted position 197 to N. Both N150K and K203R had similar effects on HAI of B/Brisbane/60/2008 197N antiserum, regardless of whether they were introduced on a 197N or 197D HA backbone (Figure S5b). However, inhibition of B/Brisbane/60/2008 by the B/Austria/1359417/2021 antiserum was only evident when N150K and K203R were introduced on a 197D backbone, highlighting the critical role of position 197. We therefore tested the remaining of mutations on a 197D B/Brisbane/60/2008 HA backbone. We confirmed that the antigenic impact (fold-change in HAI) of mutations on the 197D HA backbone was similar regardless of whether a B/Brisbane/60/2008 197N or 197D antiserum was used, with N150K having the least effect, K203R and Δ162-164 having modest effects and K136E having the greatest effect (Figure S5c). Mutations A127T, N129D, P144L, I180V and G184E on the B/Brisbane HA did not impact HAI activity of B/Brisbane/60/2008 or B/Austria anti-sera. On the other hand, K136E, Δ162-164, N150K and K203R on their own as well as in various combinations both reduced HAI by B/Brisbane/60/2008 antisera and increased HAI by B/Victoria/2023 antisera (B/Austria/1359417/2021-like) (Figure S5d-g).

Next, we assessed cluster transitions within the B/Yamagata lineage. To avoid handling of live infectious B/Yamagata viruses, due to the disappearance of B/Yamagata, the impact of mutations was tested using virus-like particles (VLP) with the relevant HA (Figure S6a). To determine the utility of this VLP-based HAI assay in antigenic characterisation, we compared HAI titers of 7 ferret antisera against 3 IBV antigens (B/Yamagata/16/1988, B/Malaysia/2506/2004, B/Phuket/3703/2013) as whole live virus or VLPs (Figure S6b). HAI titers determined by whole live virus or VLPs were highly correlated (r>0.91). We further validated the VLP-based HI assay by determining HI titers using independent batches of VLPs generated on different days (Figure S6c), as well as determining HI titers from the same set of samples on two different days (Figure S6d). In both cases, HI titers were highly correlated between experimental conditions (r>0.97). Using this system, we tested the impact of amino acid substitutions on the cluster transitions as above. Overall, the impact of mutations was more subtle than for the Ancestral or B/Victoria lineages. Inhibition of B/Yamagata/16/1988 by its homologous antiserum was not reduced but was rather increased by R162N, T168N, but not D126N. The combination of all 3 mutations, but not the individual mutations themselves increased inhibition of B/Yamagata/16/1988 by the B/Florida/04/2006 antiserum (Figure S6e). We next considered viruses represented by B/Massachusetts/02/2012 as they (i) are phylogenetically distinct, (ii) were poorly recognised by the B/Florida/04/2006 antiserum and (iii) were considered antigenically distinct for vaccine strain selection(38). However, the one mutation tested (T182A) on the B/Florida/04/2006 background did not impact inhibition by B/Florida/04/2006 nor B/Massachusetts/02/2012 ferret-antiserum (Figure S5f). Inhibition of B/Florida/04/2006 by its homologous antiserum was reduced by N150I mutation, and inhibition by the B/Phuket/3073/2013 antiserum was increased by the combined mutations N150I, N166Y, N203S (Figure S5g). This was in contrast to the impact of the 3 individual mutations, indicative of epistasis.

To identify trends in the residues involved in immune recognition by ferret antisera, we summarised the impact (absolute value of HAI fold-change) of all the mutations tested across different HA backgrounds and against various homologous and heterologous antisera (Figure 2a). When considering the maximum effect detected in our experiments positions 136, 150, 162-164, 165, 197 and 203 had an ≥8-fold effect on HAI, while other mutations (127, 129, 133, 144, 162, 168, 170, 182, 184) only had a 2-fold effect. This difference in impact was consistent when we also considered the mean effect across all titrations performed with positions 136, 150, 162-164, 165, 197 and 203 had on average >2-fold effect on HAI, while all other mutations had <2-fold impact. These residues with the greatest impact were closest to the RBS (Figure 2b).

**Figure 2.**
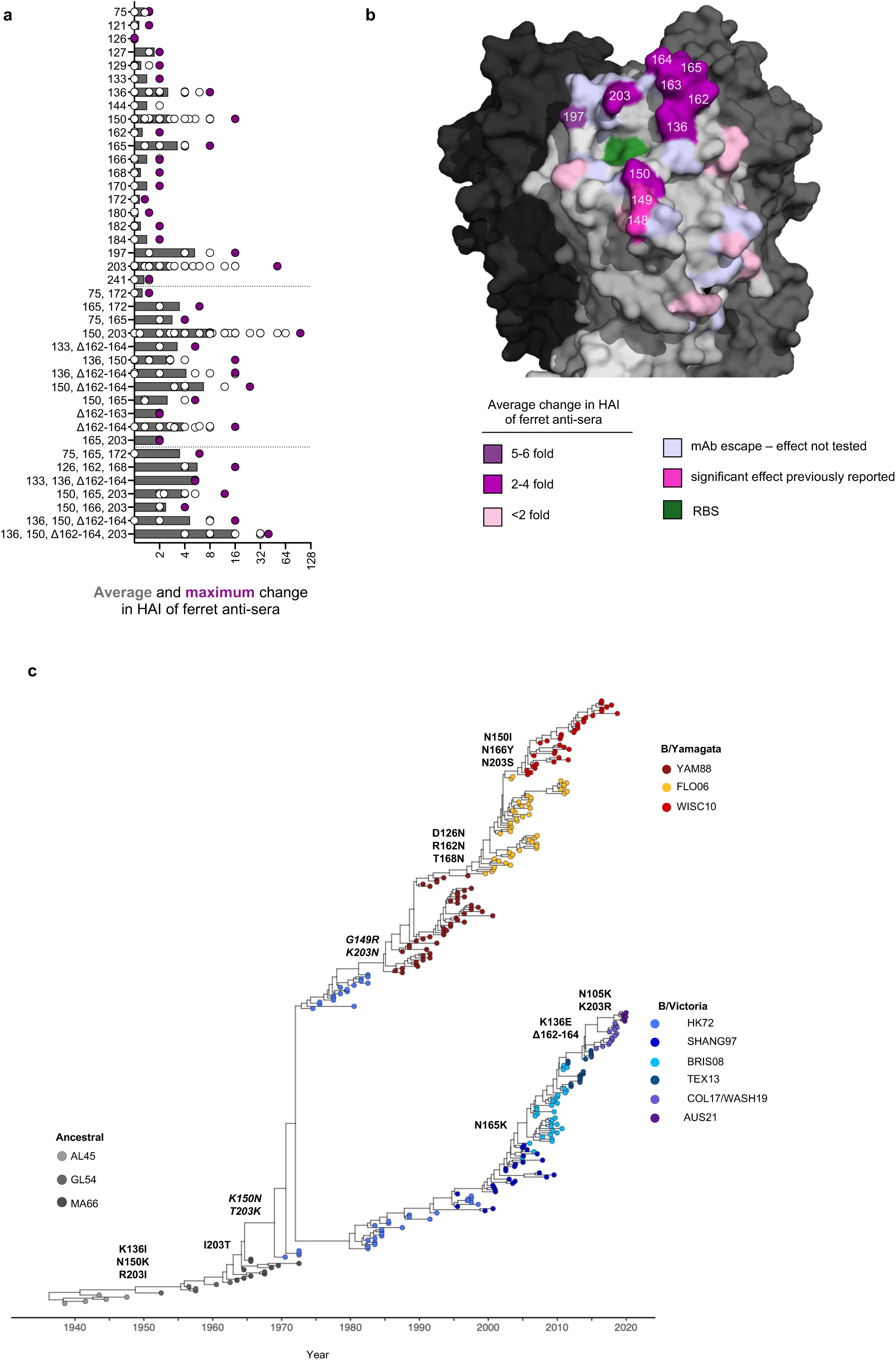
Molecular basis of IBV HA antigenic evolution. **(a)** Effect of mutations on HAI activity of ferret antisera. The average and maximum effect (absolute value of HAI change relative to wildtype) detected for each mutation tested against homologous or heterologous antisera. **(b)** Effect of mutations on ferret antisera HAI activity visualised on HA structure of B/Brisbane/60/2008 (PDB 4FQM), coloured based on the average effect detected for each mutation. Residues that have been experimentally confirmed by others for their role in IBV antigenic evolution and residues identified previously by escape mutations from mAbs but not tested here are also shown. **(c)** Temporal phylogeny of IBV coloured by the antigenic cluster and with mutations underpining antigenic evolution indicated along the trunk of the tree. Amino acid residues are numbered based on the B/Brisbane/60/2008 HA without the leader peptide. Mutations in italics were previously tested by Rosu et al (31).

Overall, from our analyses, positions 136, 150, 162-164, 165, 197 and 203 (Figure 2c) were the primary drivers of antigenic variation of IBV between 1940-2021.

### Impact of antigenic mutations on human serum

Our antigenic data so far are based on HAI using ferret antisera generated after infection with a single IBV isolate in a previously confirmed influenza seronegative animal. This does not always capture the complexity of human immunity, and the results can be influenced by the immunodominance of a given ferret antiserum(39). We thus sought to determine if mutations at key residues reduce the HAI activity of human sera. To ensure the individuals sampled had not been exposed to the mutation being tested, we focused on samples collected prior to the emergence of each mutation, or at least before their widespread circulation depending on the availability of historic samples. Future viruses that circulated after sample collection and to which these individuals had also not been exposed to were included as positive controls for immune escape. Where possible we assessed serum samples from children (1-18 years old), adults (18-60 years) or older adults (65+ years old), although we note the historic samples available to us did not allow us to directly compare the effects across age groups.

For example, to assess the impact of the N165K mutation (the major driver of the B/Malaysia/2506/2004 to B/Brisbane/60/2008 transition), we analysed serum samples collected from younger (<65 years old) and older (>65 years old) adults in 2006 (prior to 165N emerging). We also assessed samples from children (<18 years old) in 2009, the first year 165N became dominant over 165K(30), as earlier samples from children were not available. We also included mutations at the other dominant sites identified from ferret antisera (N150K, Δ162-164, K203R) and mutations identified as putatively underpinning the cluster transition (N75K, S172P) regardless of whether an effect was observed in the context of ferret anti-sera. The N165K mutation and the Δ162-164 consistently reduced HAI activity as did the N150K/K203R combination, supporting the role of these sites in antigenic recognition of the IBV HA (Figure 3a). In the context of the B/Brisbane/60/2008 HA, mutations at 129, 133, 136, 165, 184, 197 and 203 reduced HAI activity of human serum, although the average effect varied from ∼3-fold to a less than 2-fold reduction in titre (Figure 3b). In the context of the B/Florida/04/2006 HA, mutations at 150, 165, 203 and 182 resulted in reduced HAI activity of human serum, although the average effect was generally modest (2-fold reduction). The effect of the ‘future’ B/Yamagata variants (B/Massachusetts/2/2012 and B/Phuket/3703/2013) was much greater (>6-fold reduction), suggesting additional mutations influence immune escape, either directly or by epistasis. Nonetheless, these analyses collectively support the idea that substitutions at positions 136, 150, 165, 197 and 203 along with the Δ162-164 deletion have an important role on IBV immune escape from human polyclonal sera.

**Figure 3.**
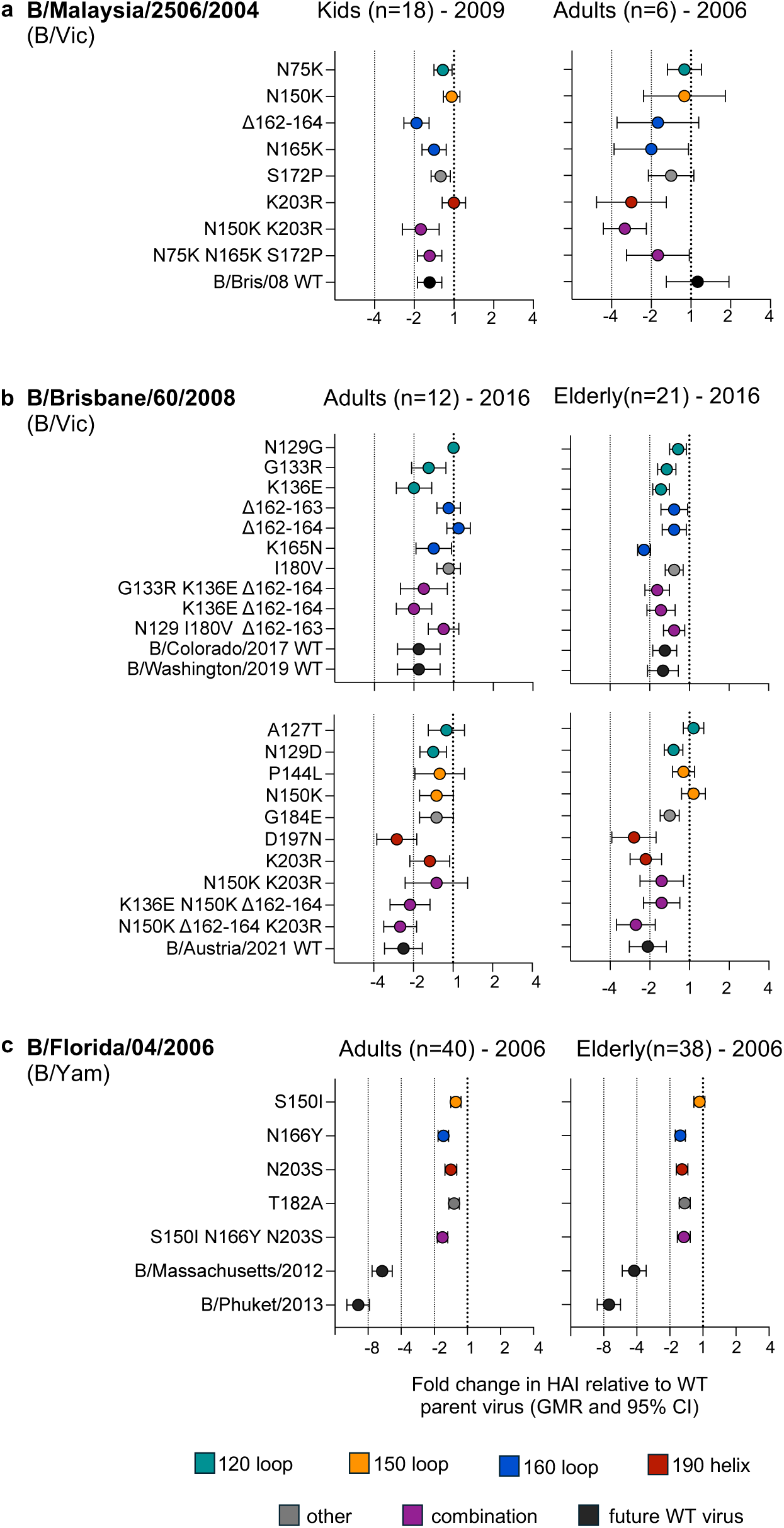
Impact of antigenic mutations on human polyclonal serum. Effect of mutations on HAI activity of human sera from different cohorts. The impact of mutations (fold change relative to wildtype) was assessed on **(a)** the B/Malaysia/2506/2004 HA (n=18 children sampled in 2009 and 6 adults sampled in 2006), **(b)** the B/Brisbane/60/2008 HA (n=12 adults and 21 older adults sampled in 2016 and **(c)** the B/Florida/04/2006 HA (n=40 adults and 38 older adults sampled in 2006).

### Molecular basis of antigenic reversion of recent B/Victoria-like viruses

Our antigenic map suggests that viruses from the B/Victoria lineage have undergone an antigenic reversion, in stark contrast to the linear path of A(H3N2) in antigenic space. Upon inspection of the raw HAI data this antigenic reversion was driven by cross-reactivity between the B/Mass/3/1966 antisera and viruses from the B/Austria/1359417/2021 cluster (Figure 3a), an observation consistent with results by Rosu et al(31). In our previous analysis we had also noted that humans born around the 1960s (but not other decades) and sampled prior to 2020 had high HAI towards the previously unencountered B/Austria/1359417/2021 strain(32), further suggestive of antigenic similarity between this virus and those from the 1960s. Alignment of key antigenic residues suggested this antigenic similarity was likely underpinned by positions 150 and/or 203 (Figure 4b). Indeed, when introduced on the B/Brisbane/2008 HA, mutations N150K and K203R increased HAI activity of both the B/Austria/2021 as well as the B/Mass/1966 antisera. This effect was also evident, and in fact greater, when N150K and K203R were combined with K136E and Δ162-164, which are not present in B/Mass/1966 (Figure 4c). These data demonstrate that the recycling of IBV HA mutations can lead to the re-emergence of past antigenic phenotypes.

**Figure 4.**
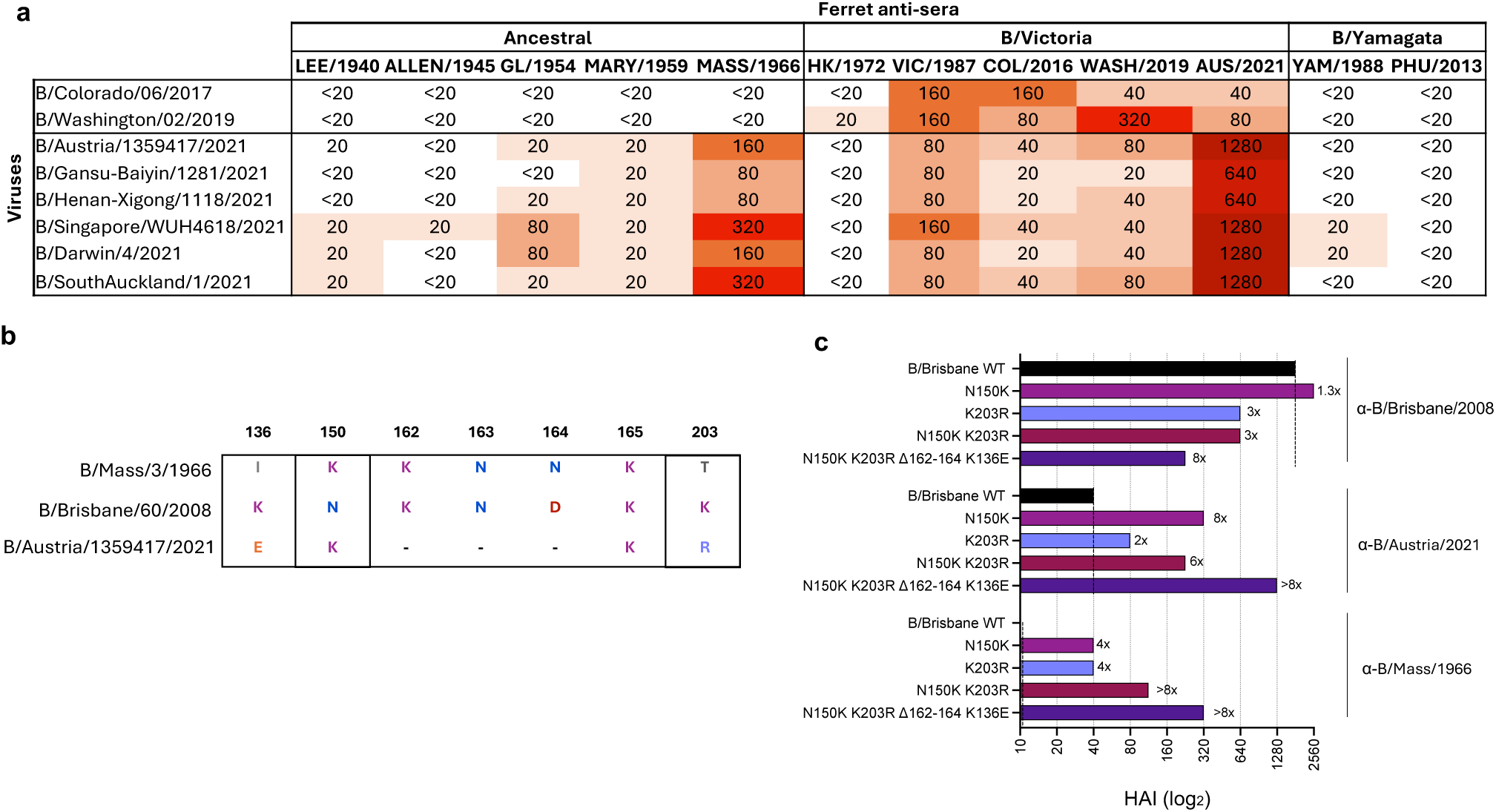
Molecular basis of IBV HA antigenic reversion. **(a)** Heatmap of HAI data of 12 ferret antisera against 8 IBV isolates. **(b)** Sequence alignment of key antigenic residues. **(c)** Impact of mutations on the B/Brisbane HA on HAI activity of B/Austria/2021 and B/Mass/1966 specific ferret antisera. The fold change relative to the HAI titre against WT B/Brisbane is shown (e.g. 2x) and the dashed vertical lines indicate the reference comparison for each antiserum. All antisera had a homologous titre of >960.

### Epistatic interactions shape the impact of mutation recycling

Given the impact of mutation recycling (Figure 4), we then assessed amino acid usage in the 10 key antigenic residues (Figure 2c) across different clusters from the Ancestral and B/Victoria lineage (Figure 5a). As changes at position 197 may be due to egg passage, we did not consider it in this analysis. Across the other 9 residues, we noted that certain amino acids were frequently reused at positions 136, 150, 162-164, 165 and 203 (Figure 5b).

**Figure 5.**
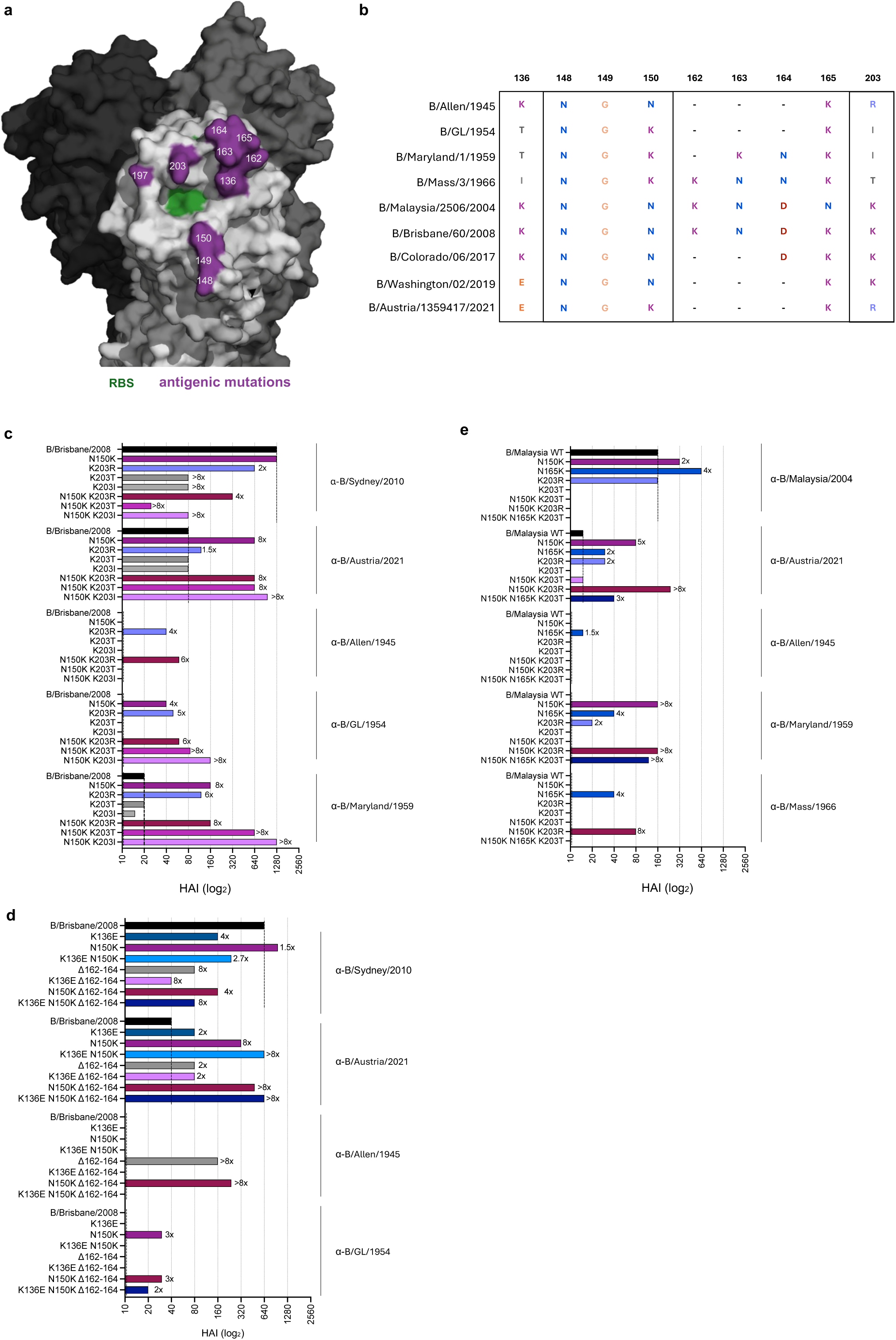
Epistatic interactions shape the impact of mutation recycling. **(a)** Key antigenic residues of the IBV HA of B/Brisbane/60/2008 (PDB 4FQM). **(b)** Sequence alignment of key antigenic residues across IBV isolates from the Ancestral and B/Victoria lineage. **(c)** Impact of mutations at 150 and 203 on the B/Brisbane HA on HAI activity of ferret antisera. All antisera had a homologous titre of >640. **(d)** Impact of mutations at 136 and 150 and the Δ162-164 deletion on the B/Brisbane HA on HAI activity of ferret antisera. All antisera had a homologous titre of >640. **(e)** Impact of mutations at 150, 165 and 203 on the B/Malaysia HA on HAI activity of ferret antisera. All antisera had a homologous titre of >160. **(c-e)** The fold change relative to the HAI titre against WT B/Brisbane (c-d) or B/Malaysia (d) is shown (e.g. 2x) and the dashed vertical lines indicate the reference comparison for each antiserum.

To understand the impact of this amino acid recycling, we firstly assessed substitutions at 203 on the B/Brisbane/2008 HA (Figure 5c). All 203 substitutions tested reduced HAI activity of the B/Sydney/508/2010 anti-serum (B/Brisbane/60/2008-like) and did not affect HAI activity of the B/Austria/1359417/2021 anti-serum, with or without N150K (Figure 5c). Only K203R increased HAI activity of the B/Allen/1945 antiserum, consistent with the presence of 203R in B/Allen/1945, even when combined with N150K for which B/Allen/1945 has an N. When an T or I were introduced at 203 on their own, they did not impact HAI activity of the B/GL/1954 or B/Maryland/1959 antisera. However, when combined with the N150K mutation, they increased HAI activity of both antisera by 2-8-fold times over the impact of N150K alone, indicative of epistatic interactions. This exemplifies how the effects of some antigenic mutations depend on the presence of other mutations.

Next, we assessed the impact of mutation recycling in the 160-loop, specifically the Δ162-164 deletion, and mutations at 136, which is structurally adjacent to residues 162-165 (Figure 5a). Both, the B/Syndey/508/2010 (B/Brisbane/60/2008-like) and the B/Austria/1359417/2021 antisera were highly sensitive to changes at these sites on the B/Brisbane/60/2008 HA. Importantly, Δ162-164 on the B/Brisbane/2008 HA increased HAI activity of the B/Allen/1945 antiserum, which shares Δ162-164 (Figure 5b). However, when combined with K136E, this completely abrogated the impact of Δ162-164, despite K136E having no effect on its own. This was not the case for combining N150K and Δ162-164 (Figure 5c), indicating the specific nature of these epistatic interactions. We also noted that Δ162-164 did not increase HAI activity of the B/GL/1954 anti-serum (Figure 5c), which also contains shares Δ162-164, further suggesting that these effects are context specific. These results overall exemplify how the effects of some antigenic mutations depend on the absence of other mutations.

Finally, we assessed whether the introduction of mutations at positions 150, 165 and 203 on the B/Malaysia/2506/2004 HA resulted in recycling of antigenic phenotypes as they did on the HA of B/Brisbane/60/2008(Figure 5d). The B/Malaysia/2506/2004 anti-serum was highly sensitive to the combination of mutations at positions 150 and 203. The N150K mutation increased HAI activity of the B/Austria/1359417/2021 antisera, and this was amplified by the K203R, but not K203T mutations. The HAI activity of the B/Allen/1945 antiserum was not impacted by any of the mutations tested, in contrast to the effect of these mutations for the same serum in the context of the B/Brisbane/2008 HA (Figure 5c). The HAI activity of the B/Maryland/1959 antiserum increased with the N150K mutation, but not by the K203T mutation. In contrast, the HAI activity of the B/Mass/1966 antiserum was not impacted by the N150K or K203R/T on their own but increased by the combination of N150K and K203R. The N165K mutation on its own increased HAI activity of all antisera to variable extents (Figure 5d). These results further demonstrate the epistatic interactions between these antigenic residues and how they can be highly variable depending on the presence or absence of other mutations.

Overall, our data clearly demonstrate that residues 136, 150, 162-165, and 203 form a highly interconnected epistatic network which modulates their antigenic effects.

### Epistatic interactions rescue deleterious effects of antigenic mutations

We next considered the potential role epistasis plays in balancing viral fitness and immune escape. We assessed IBV replication in air-liquid interface (ALI) cultures of human airway epithelial cells. Human nasal epithelial cells (hNECs) were obtained from nasal brushings and used to generate well-differentiated cultures at an ALI, while BCi-NS1.1(40) an immortalized bronchial basal epithelial cell line, was used to generate differentiated large airway epithelial cells (LAECs) cultures, and hSABCi-NS1.1(41) an immortalized small-airway basal epithelial cell line, was used to generate small-airway epithelial cells SAECs cultures, as previously described(42). All three types of ALI cultures supported IBV replication, albeit with differential kinetics (Figure S7a). Replication kinetics were mostly identical for a wild-type B/Malaysia/2506/2004 derived from a clinical isolate and a recombinant virus rescued by reverse genetics, with only a small difference in the later stages of infection in hNECs (Figure S7a).

We firstly compared viral replication of B/Malaysia/2506/2004 viruses with the single or combined N75K, N165K, and S172P mutations in hNECs and LAECs (Figure 6a-b) (Table S2). None of these mutations substantially affected viral replication kinetics compared to WT viruses. We also assessed replication *in vivo* and did not detect any growth defects of the mutated viruses in the upper (nasal tissue) or lower (lung tissue) respiratory tract of C57BL/6 mice on day 3 post infection (peak viral load)(43)(Figure S7b), indicating that mutations at 75, 165 and 172 on B/Malaysia/2506/2004 had no major effect on viral fitness.

**Figure 6.**
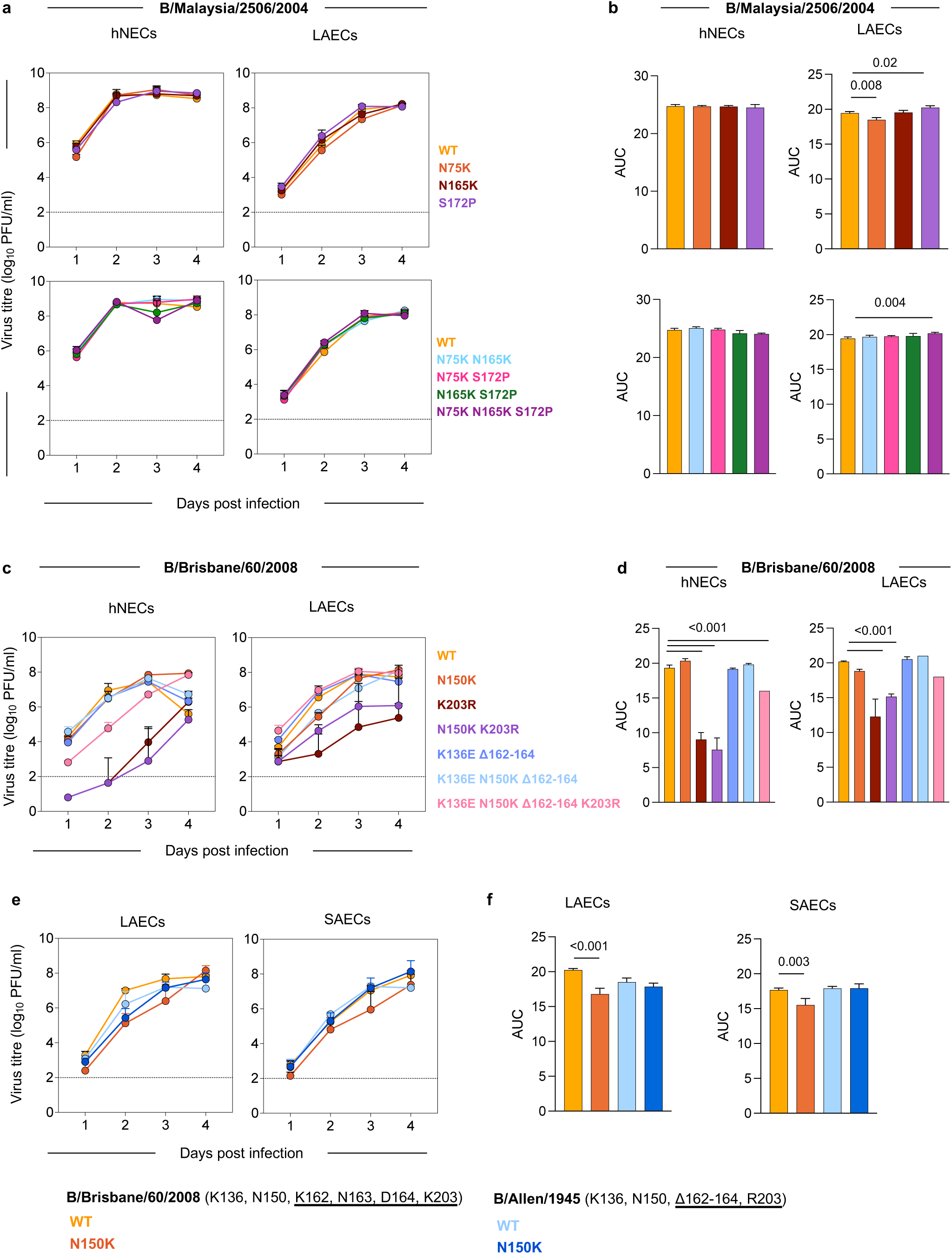
Epistatic interactions rescue deleterious effects of antigenic mutations. (a-b) Impact of mutations on the B/Malaysia/2506/2004 HA, shown as virus titres (PFU/mL) across time (a) and area under the curve (b). Mean and standard deviation from triplicates are shown. **(c-d)** Impact of mutations on the B/Brisbane/60/2008 HA, shown as virus titres (PFU/mL) across time (c) and area under the curve (d). Mean and standard deviation from triplicates are shown. Data representative of two experiments, except for the K136E, 150K, Δ162-164, K203R mutant which was not available in the first experiment. **(e-f)** Impact of N150K mutation on the B/Brisbane/60/2008 and B/Allen/1945 HAs, shown as virus titres (PFU/mL) across time (E) and area under the curve (F). Mean and standard deviation from triplicates are shown. Throughout the figure, ALI differentiated human airway nasal epithelial cells (hNECs), large-airway epithelial cells (LAECs) or small-airway epithelial cells (SAECs) were inoculated at a multiplicity of infection (MOI) of 0.01, sampled daily after infection and viral titres were determined by plaque assay. Dotted lines indicate the limit of detection. Statistical significance for viral titres was determined by two-way ANOVA followed by Dunnett’s multiple comparison test and is summarised in table S2. Statistical significance for AUC was determined by a one-way ANOVA comparing each mutant the WT with Dunnett’s correction for multiple comparisons.

We next compared viral replication of B/Brisbane/60/2008 viruses with single or combined mutations at key antigenic positions (Table S2). The K203R mutation alone or in combination with N150K had a strong deleterious effect on viral fitness, resulting in delayed and overall reduced viral replication (Figure 6c-d). The N150K mutation only had a small effect on viral replication on its own in LAECs. Combining the N150K/K203R mutation together with K136E and Δ162-164 significantly restored viral fitness. These phenotypes were consistent in both hNECs and LAECs. We note that the K136E/Δ162-164/N150K/K203R mutant did not reach WT B/Brisbane/60/2008 replication levels, suggesting that some of the additional mutations in that cluster transition (A127T, N129D, P144L, I180V, G184E) likely further compensate the deleterious effects of N150K/K203R mutations. As the K136E/Δ162-164 mutations preceded the N150K/K203R mutants in the IBV HA phylogeny (Figure 2b), our data indicate that these mutations generated a permissive backbone on which the otherwise deleterious N150K/K203R mutations could emerge with limited fitness cost.

To further understand the context-dependent effects of immune escape mutations on viral fitness, we compared the impact of the N150K mutation on B/Allen/1945 and B/Brisbane/60/2008, since both strains naturally acquired this mutation which altered their antigenic phenotype. These two strains also differ in some other key positions like 203 (R in B/Allen/1945, K in B/Brisbane/60/2008) and the 160 loop (Δ162-164 in B/Allen/1945 vs K162 N163 D164 in B/Brisbane/60/2008), but both have a K at position 136. The N150K mutation in B/Brisbane/60/2008 but not B/Allen/1945 resulted in reduced replication levels LAECs as well as SAECs (Fig 6e-f). These data highlight the context dependent effects of mutations on viral fitness and the importance of epistatic interactions in shaping viral fitness.

## Discussion

Our study of IBV HA evolution over 8 decades identifies critical similarities and differences with the evolution of A(H3N2) and demonstrates the critical role of epistasis in balancing antigenic novelty with viral fitness. Our observations demonstrate how epistatic interactions may (i) increase the number of possible antigenic phenotypes despite the small number of residues that can facilitate antigenic change, and (ii) increase accessibility to such antigenic phenotypes with limited fitness cost. In turn, these findings explain how antigenic evolution can be facilitated by a small number of residues, even over 8 decades of circulation in humans, while maintaining fitness. This emphasizes the importance of further understanding the molecular basis of viral evolution in order to develop strategies that can effectively predict and combat it.

In addition to demonstrating the importance of epistasis, we provide the most comprehensive characterisation of the IBV antigenic space to date. Despite the initial identification of IBV HA antigenic residues using mAbs more than 4 decades ago(44, 45), the antigenic space of the IBV HA had only been partially mapped(31, 46–51). Importantly, the amino acid residues that drive IBV antigenic evolution in nature had not been experimentally validated, with the exception of the residues that resulted in the emergence of the B/Victoria and B/Yamagata lineages (positions 148, 149, 150 and 203)(31). Our work, combining ferret and human sera, demonstrates that positions 136, 150, 162-165, 197 and 203 are important determinants of the observed antigenic evolution between 1940-2021, with positions 126, 129, 133 and 168 also playing a potential role. Overall, our data in combination with Rosu et al(31), suggest that 10 amino acid residues (136, 148-150, 162-165, 197 and 203) underpin the antigenic evolution of the IBV HA across the 3 antigenic lineages and 12 antigenic clusters over the last 8 decades. Accordingly, these residues should be considered in influenza virus surveillance and vaccine strain selection. It is pertinent to note that additional antigenic clusters and mutations might have antigenic effects not assessed here. For instance, the V1A.3a.1 phylogenetic cluster that briefly circulated in China in 2021 (52), and was thus not captured in our dataset, contains the mutations V220M and P241Q, which we have not assessed, and appears antigenically distinct from and the V1A.3a.2 (B/Austria/1359417/2021) cluster(49, 50). Similarly, the recently emerged C.3.1 cluster, which contains D197N, appears antigenically distinct from B/Austria/1359417/2021(53, 54). Therefore, antigenic states other than those observed between 1940-2021 are likely possible for the IBV HA, and their potential molecular basis needs to be determined.

The recycling of antigenic phenotypes and the resulting non-linear antigenic path of the IBV HA across antigenic space is another stark contrast with A(H3N2). Whether it represents a limitation of antigenic options for the IBV HA is difficult to ascertain at present as it has been a single, and potentially stochastic, occurrence. We note that mutations have also been recycled in A(H3N2)(55), but this has not, so far, resulted in the re-emergence of past A(H3N2) antigenic phenotypes. Understanding the potential limits of the IBV antigenic space and the underlying molecular basis will inform on the feasibility of universally protective vaccines.

IBV uses a relatively small number of residues with even single mutations mediating antigenic cluster transitions, similar to different IAV subtypes(6, 9, 56–58). For both IBV and A(H3N2), mutations often result in biochemical changes (e.g. charge changes). In contrast to A(H3N2), however, we found that IBV additionally utilises insertions and deletions (indels), specifically in the 160-loop. While both substitutions and indels likely impact antibody binding directly, they may also modulate receptor avidity resulting in differential sensitivity to inhibition(59), which we have not experimentally assessed for the IBV HA. In addition, we confirmed the previously reported antigenic impact of glycosylation changes at position 197-199 of the IBV HA(34, 35), consistent with the antigenic impact of glycosylation in A(H3N2) evolution(60). It is worth noting that 160-loop indels immediately precede a putative glycosylation site (N166, K167, T168) that is conserved across Ancestral and B/Victoria viruses. It is possible that the antigenic impact of indels is repositioning of the glycans. Similarly, the T168N mutation that occurred in the B/Yamagata lineage both disrupts the N166-K167-T168 motif and introduces a glycosylation site at N168-K169-T170, thus shifting the position of the glycan. Given these observations, the impact of mutations at the glycosylation site 197-199, and the emergence of the C.3.1 cluster with a re-introduced 197-199 glycosylation site (53), it remains pertinent to perform additional structural and biochemical studies to fully understand the impact of glycosylation changes in IBV antigenic drift as well as viral fitness.

The diversification of the IBV HA into two antigenic lineages is a significant difference to A(H3N2), although whether IBV generally has a greater propensity for such speciation events is unclear since this has been a single occurrence. Interestingly, while 2/3 mutations that gave rise to the B/Yamagata lineage (N148S, G149R)(31) have remained relatively conserved, our data showed that position 203 evolved further in the early 2010s (N203S). Similarly, we show that mutations at positions 150 and 203, which defined the emergence of the B/Victoria lineage(31), also occurred in the 1950s and in the early 2020s but did not result in the establishment of a new co-circulating antigenic lineage. Therefore, while mutations at these residues appear antigenically necessary, they are not sufficient for the generation of novel antigenic lineages that can co-circulate with existing lineages. This further highlights the impact epistasis has in determining the outcome of mutations at these key sites. A greater understanding of the molecular basis of B/Yamagata emergence is needed to understand the likelihood of a similar event occurring again. It is also tempting to speculate on the potential role of epistasis in the eventual extinction of B/Yamagata. While we found that epistasis can diversify antigenic effects and rescue viral fitness for IBV, there’s evidence that epistasis can also limit the evolution of A(H3N2) via entrenchment(12, 15). The degree to which entrenchment has directed the evolution of IBV deserves further attention.

While our study has several strengths, including the analysis of >400 viruses representing 8 decades of viral evolution, the experimental assessment of >60 mutant viruses and the use of ALI cultures to assess viral replication, it also has limitations. Firstly, some of the viruses analysed were isolated in eggs and carry the egg-adaptive loss of the glycosylation site at position 197-199 of the IBV HA. We have, however, confirmed that this does not impact the primary conclusions of our study (Figure S1d,c and S5b,c). Secondly, we have not tested the impact of all possible mutations or their permutations, and the antigenic basis of some cluster transitions was not entirely clear. Given the extent of epistasis observed among the mutations tested, and as discussed above, it is possible that additional mutations, on their own or in combinations, modulate the antigenic and fitness phenotypes of the IBV HA. Additionally, our antigenic analyses are based on HAI assays, and live virus microneutralization assays may yield epidemiologically relevant insights into immune escape from antibodies that do not mediate HAI. Furthermore, we have not determined the molecular mechanisms behind the observed epistasis for antigenic phenotypes or viral fitness, which may be pleiotropic(12). Finally, more systematic approaches to map epistasis can be employed to identify additional pairwise or higher-order interactions across the HA as well as between the HA and NA, which we have not considered here. Despite these limitations, our data clearly demonstrate that mutations at a small number of RBS-proximal residues result in large antigenic changes of the IBV HA, and the effect these residues have on antigenicity and replicative fitness are strongly influenced by epistasis.

A comprehensive understanding of the processes that underpin antigenic evolution and developing a unifying framework of viral evolution can facilitate the development of effective strategies to predict and combat viral evolution. To date, this has primarily relied on the study of influenza A(H3N2) viruses. Our comprehensive antigenic and experimental characterisation of IBV HA evolution over >8 decades highlights both differences and similarities with A(H3N2) and provide novel insights for the development of viral evolution frameworks. There is also increasing interest in integrating genetic and antigenic data for the development of tools that can predict phenotypes from sequence data(61–64). Their development has similarly relied on seminal papers mapping the antigenic evolution of A(H3N2)(6, 9). Our sequence, antigenic and viral fitness data can be an additional valuable resource to further develop such tools. These tools must however capture, and ideally forecast, the often unpredictable effects of epistasis.

## Methods

### Viruses and serum samples

Details of influenza B isolates used in this study are provided in Table S3. Egg-grown isolates were propagated in 10-12 day old embryonated chicken in accordance with guidelines set by the University of Melbourne Animal Ethics Committee (ethics approval number 10448-2015136 and 27270). Cell-grown isolates were propagated in Madin Darby Canine Kidney (MDCK) cells in the presence of TPCK-treated trypsin. Viral RNA was extracted from clarified supernatants, and HA segments were amplified and sequenced by Sanger sequencing; sequences were aligned using the multiple alignment using fast Fourier transform (MAFFT) algorithm implemented in MegAlign Pro 13 (DNASTAR Lasergene 13).

Ferret antisera (Table S3) were provided by the WHO Collaborating Centre for Reference and Research on Influenza (WHO CCRRI, Melbourne) and were generated in accordance with guidelines set by the University of Melbourne Animal Ethics Committee (ethics approval number (10423-2015104 and 27365) and by the CSL Animal Ethics Committee (ethics approval number 2040-1). Human serum samples were collected from participants who provided written informed consent in accordance with the Declaration of Helsinki and as approved by the relevant ethics committees. Adult human sera were collected under study protocols approved by the University of Melbourne Human Research Ethics Committee (1443420) or provided by the WHOCCRRI in Melbourne as outlined in the terms of reference of the WHO Global Influenza Surveillance and Response System. Samples from children aged 1–18 years in 2009 were previously collected as part of a clinical trial (NCT00959049)(65).

### HAI assays

HAI assays were performed as previously described(32) and according to the WHO Global Influenza Surveillance Network protocols(66), with the exception that volumes were reduced to 25 μL of sera, virus (4 HA units) and 1% turkey erythrocytes (0.33% final concentration). Ferret and human sera were treated with receptor destroying enzyme (Denka Sieken) and adsorbed with 5% erythrocytes prior to testing. Samples were tested over two-fold serial dilutions from 1:20 to 1:20,480. Each assay included ferret antisera and/or monoclonal antibodies(67) with known reactivity patterns as positive controls, and PBS as a negative control.

### Antigenic and sequence data

We assembled a dataset of HAI measurements for 287 IBV isolates from 1940-2021. The core dataset comprised lineage-specific HAI measurements generated at the WHOCCRRI, Melbourne, in which B/Yamagata-specific antisera were tested against B/Yamagata isolates, and B/Victoria-specific antisera tested against B/Victoria isolates from 2002–2013. These were previously reported by Dhanasekaran et al (46). We expanded this dataset by re-testing the same ferret antisera against additional virus isolates and by testing the same original viral isolates against additional ferret antisera at WHOCCRRI using identical protocols. We further incorporated HAI measurements reported by Rosu et al (31) for 64 IBV viruses tested against 16 ferret antisera from 1958–2000. The final combined dataset comprised 287 viruses and 66 ferret antisera, with 8,008 completed serum-virus titrations representing 43.3% of all possible combinations (Table S4). Viral HA gene sequences (table S3) were obtained either from virus stocks generated in this study or downloaded from GISAID (68, 69). Amino acid numbering followed the B/Brisbane/60/2008 HA sequence (GenBank accession FJ766840), which lacks deletions in the 160-loop, with residue 1 corresponding to the aspartic acid in the DRIC motif following the signal peptide. This numbering scheme was applied to all analyses and is consistent with WHO reports(36), NextStrain (30), and previous studies of IBV HA (31).

### Phylogenetics incorporating antigenic evolution

Time-scaled phylogenetic trees were inferred from 405 HA gene sequences using BEAST v1.10.1(70) under a general time reversible (GTR) + Γ4 nucleotide substitution model, an uncorrelated lognormal relaxed molecular clock, and a Gaussian Markov random-field (GMRF) Bayesian Skyride coalescent prior. MCMC chains were run at least twice independently for 50 million steps, sampling every 5,000 steps, and discarding the first 10% as burn-in. From the combined posterior, 4,500 time-scaled trees were randomly sampled to generate an empirical tree distribution used in downstream antigenic analyses.

Antigenic evolution was modelled as a continuous diffusion process over the shared viral phylogeny using a two-dimensional Bayesian multidimensional scaling (BMDS) model(47) applied to 287 HA sequences with corresponding HAI data, comprising 8,008 virus– antiserum pairs. The model jointly estimated antigenic locations for viruses and antisera, while accounting for virus avidity and serum potency, allowing antigenic phenotypes to evolve along phylogenetic branches. BMDS MCMC analyses were run at least twice independently for 500 million steps, sampling every 200,000 steps, with 10% burn-in. Convergence and mixing were assessed using Tracer v1.7.1(71), ensuring effective sample size greater than 200 for all parameters.

Codon-based selection was performed using HyPhy (72)v2.5.8 on HA codon alignments. We applied analyses aBSREL(73), BGM(74), BUSTED(75), FEL(76), FUBAR(77), MEME(78), SLAC(76), FADE(76) to (i) the complete dataset, (ii) sequences sampled prior to divergence of the B/Victoria and B/Yamagata lineages, and (iii) lineage-specific datasets for B/Victoria and B/Yamagata viruses. Sites were considered to show evidence of positive selection when supported by at least one method at default significance thresholds (for example, P<0.05 or posterior probability ≥ 0.9, as appropriate for each model).

Antigenic clusters were identified using K-means clustering applied to the inferred BMDS antigenic coordinates and sampling time of tip viruses. Variables were standardized prior to analysis, and the optimal number of clusters was determined using the elbow and silhouette methods, resulting in a 12-cluster solution. Cluster assignments were subsequently examined in the context of the antigenic map and phylogeny.

### Reverse genetics for the generation recombinant IBVs

Reverse genetics systems for influenza B/Brisbane/60/2008 (pDP plasmid backbone) and B/Malaysia/2506/2004 (pDZ plasmid backbone) were kindly provided by D. Perez and P. Palese and used to generate reverse genetic viruses, as described before(22, 37). Briefly, 8 bidirectional plasmids were transfected into 293T cells (each 1 μg/plasmid), using FUGENE 6 transfection reagent. 24 hrs post transfection, cells were co-cultured for 2-3 days with MDCKs in the presence of TPCK-treated trypsin to allow virus replication. Clarified co-culture supernatants were obtained by centrifugation and used to inoculate MDCK cells for the generation of virus stocks as described above. For B/Allen/1945 and B/GL/1954 reverse genetic viruses, the HA and NA genes from B/Allen/1945 and B/GL/1954 were cloned from virus stocks into a bidirectional plasmid and used to generate recombinant viruses with the surface (HA, NA) proteins from B/Allen/1945 or B/GL/1954 and the internal plasmids from B/Brisbane/60/2008. Amino acid substitutions or deletions were introduced into the different HA genes either by using the Q5 site-directed mutagenesis kit (NEB), overlap extension PCR with mutagenesis primers or synthesized commercially as geneblocks (Thermo Fisher). The presence of correct mutations and absence of undesired mutations was confirmed by Sanger sequencing of the HA gene of the plasmid and the virus stock.

### Virus-like particles

293T cells were cultured in DMEM, supplemented with 1% FCS, 1mM sodium pyruvate (all Thermo Fisher) and PSG. pCMV and pDZ HA plasmids were co-transfected with pHDM-Hgm2 and pCMV B/Florida/2006 NA into 293Ts using Lipofectamine 3000 (Thermo Fisher), following the manufacturer’s instructions. 3 days post transfection, supernatant was harvested and clarified supernatants were obtained by centrifugation. VLPs were diluted to 4 HA units per 25μL and used in HAI assays as described above.

### ALI cultures and *in vitro* virus replication

ALI cultures from BCi-NS1.1 and hSABCi-NS1.1 were prepared as previously described(42). Briefly, cells were seeded in transwell plates (6.5 mm, Corning) and cultured in Pneumacult-Ex Plus medium. Upon reaching confluence, the apical chamber was exposed to air, while the basolateral chamber was replaced with Pneumacult ALI maintenance medium supplemented with 4mg/mL Heparin, 480 ng/mL Hydrocortisone and 50U/ml Penicillin and 50mg/ml Streptomycin. Media was changed 3 times per week. Cells were cultured for a minimum of 21 days at 37°C which allowed the formation of differentiated polarized cultures. For nasal cell ALI cultures, healthy individuals were recruited for this study under ethics approval HREC/35132. Nasal epithelial cells were sampled and cultured as described previously(79, 80).

For infection, the apical chamber containing the differentiated cells was washed once with warm PBS (with Ca^2+^/ Mg^2+^) and inoculated at 35°C with IBV at an MOI of 0.01 in a total volume of 100μL. After 2 hrs, the inoculum was removed, and cells were washed three times with PBS and rested at 35°C. To assess virus replication and growth, cells were incubated daily with 200μL PBS for 10 min at 35°C. Samples were collected for 4 days post infection, stored at-80°C, and viral titres were determined by plaque assay on MDCKs as described(43, 81).

### Mouse infections and *in vivo* virus replication

Male C57BL/6L mice aged 6–12 weeks were obtained from the Biological Research Facility in the Department of Microbiology and Immunology at the University of Melbourne (Melbourne, Victoria). All animal work was conducted in accordance with guidelines set by the University of Melbourne Animal Ethics Committee (ethics approval number 21799). Mice were intranasally infected with 10^3^ tissue culture infectious dose 50 (TCID_50_) of IBV diluted in 50 µL of phosphate-buffered saline (PBS) under isoflurane anaesthesia. Three days post infection, animals were culled, and lung and nasal turbinates were isolated. The tissues were homogenized and clarified supernatants were obtained by centrifugation. Lung and nasal tissue samples were stored at-80°C and viral titres were determined by plaque assay as described(43, 81).

## Statistical analysis

Viral titres for each mutant were compared to the WT virus at each time-point using a two-way ANOVA, with Dunnett’s correction for multiple comparisons. AUCs were compared for each mutant were compared to the WT virus at each time-point using a one-way ANOVA, with Dunnett’s correction for multiple comparisons. Correlations were assessed using the Spearman correlation co-efficient. All statistical analyses were performed in Prism 11 (GraphPad).

## Data availability

The raw HI datasets used for antigenic cartography, together with HA sequences, associated metadata, GISAID accession numbers, and acknowledgements, are provided in the Supplementary Information files and on Github. Code and data used for all phylogenetic analysis, Bayesian MDS, selection analyses and figure generation are available at https://github.com/vjlab/IBV-HA-evo.

## Acknowledgements

The work has been generously supported by the Morningside Foundation (M.K), the Australian National Health and Medical Research Council (M.K. S.J.K, A.K.W), the NIAID Centers of Excellence for Influenza Research and Response (CEIRR; contract 75N93021C00016) (V.D.), the Research Grants Council of the Hong Kong SAR Theme-based Research Scheme (project T11-712/19-N) (V.D.), and the Research Grants Council of the Hong Kong Special Administrative Region, China (Project No. HKU PDFS2425-7S01) (R.X). We are grateful to the CSL investigators who originally initiated the NCT00959049 trial; Ron Fouchier for sharing viruses; and to Ronald Crystal for sharing the BCi-NS1.1 and hSABCi-NS1.1cells. The WHOCCRRI is supported by the Australian Government Department of Health. The funders had no role in study design, data collection and analysis, decision to publish or preparation of the paper. We gratefully acknowledge all data contributors, i.e., the Authors and their Originating laboratories responsible for obtaining the specimens, and their Submitting laboratories for generating the genetic sequence and metadata and sharing via the GISAID Initiative, on which this research is based. The computations were performed using research computing facilities offered by Information Technology Services, the University of Hong Kong. For the purposes of open access, the author has applied a CC BY public copyright licence to any Author Accepted Manuscript version arising from this submission.

## Author contributions

S. U. S, E.R., M.A, N.S., YM.D., M.G, and M.K. performed experiments and data collection. L. S. U. S, R. X., S.H, V. D. and M.K. analysed data. R.S, K. L., S. R., S. J. K, A K. W, I. G. B. provided critical samples, reagents and resources. L. S. U. S, R. X., M. J. G., K. S., S. J. K, A K. W, V. D., M.K. contributed to the development of ideas and/or methodology. L. S. U. S, R. X., V. D. and M. K drafted the manuscript. All authors reviewed the final of version manuscript.

## Competing interest

M.K. has acted as a consultant for Sanofi group of companies. S.R. and K.L. are employees of Seqirus, an influenza vaccine manufacturer. I.G.B. has shares in an influenza-vaccine-producing company. The other authors declare no competing interests.

