## Supplementary figures for "Epistasis facilitates the long-term antigenic evolution of the influenza B virus hemagglutinin"

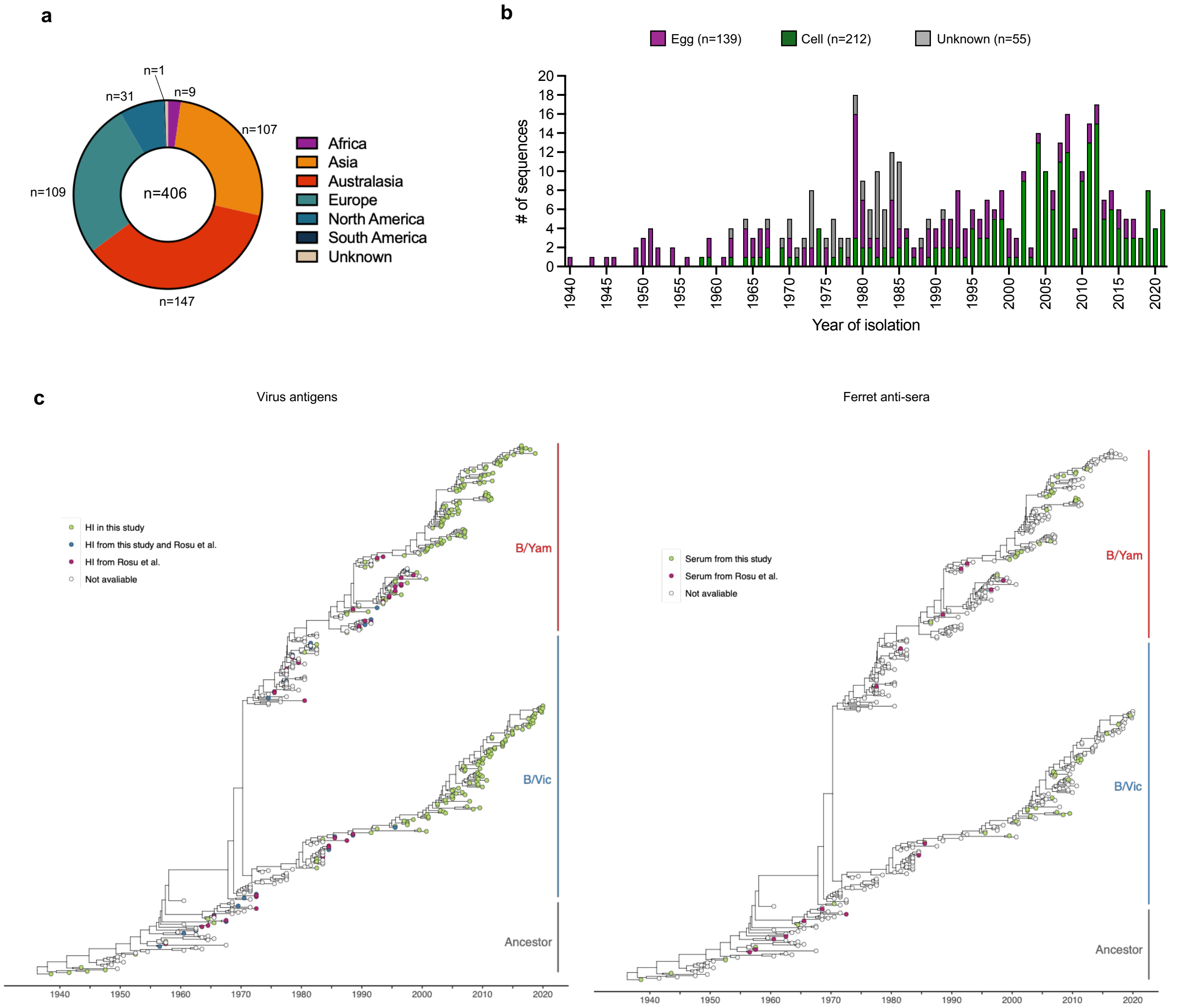


**Figure S1. Overview of IBV HA dataset and antigenic characterisation. (a)** Geographic distribution of IBV HA sequences. **(b)** Temporal distribution and egg or cell passage of HA sequences. **(c)** Viruses and antisera that were included in antigenic cartography of IBV HA highlighted in the temporal phylogeny of IBV.

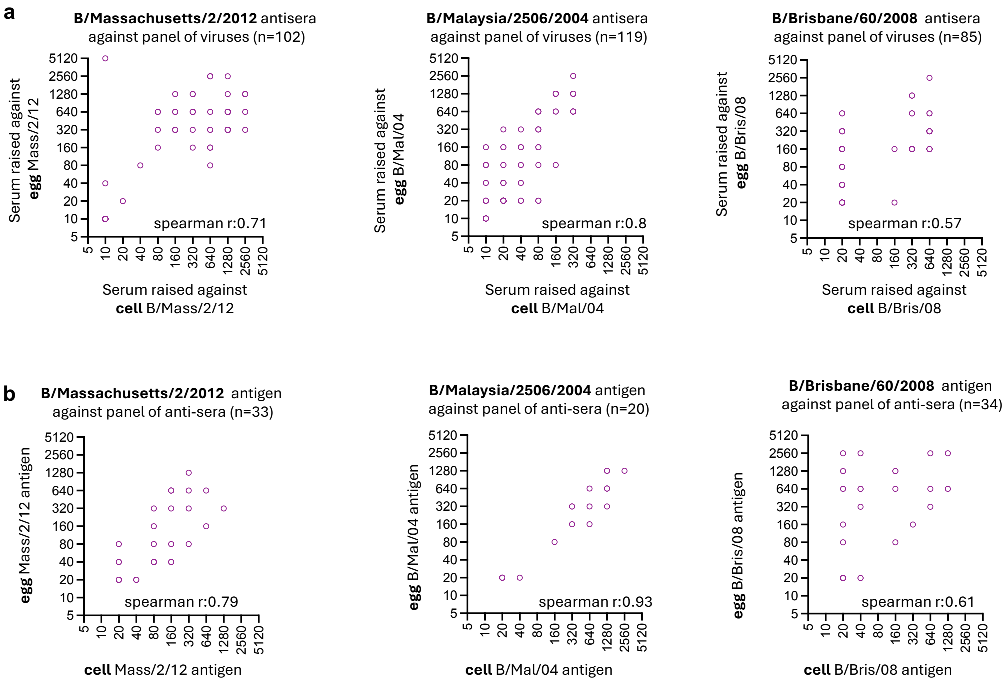


**Figure S2. Impact of egg passage. (a)** Comparison of HAI titres from ferret anti-sera raised against cell or egg-grown viruses and titrated against a panel of IBV isolates. **(b)** Comparison of HAI titres from a panel of ferret anti-sera titrated against cell or egg-grown viruses.

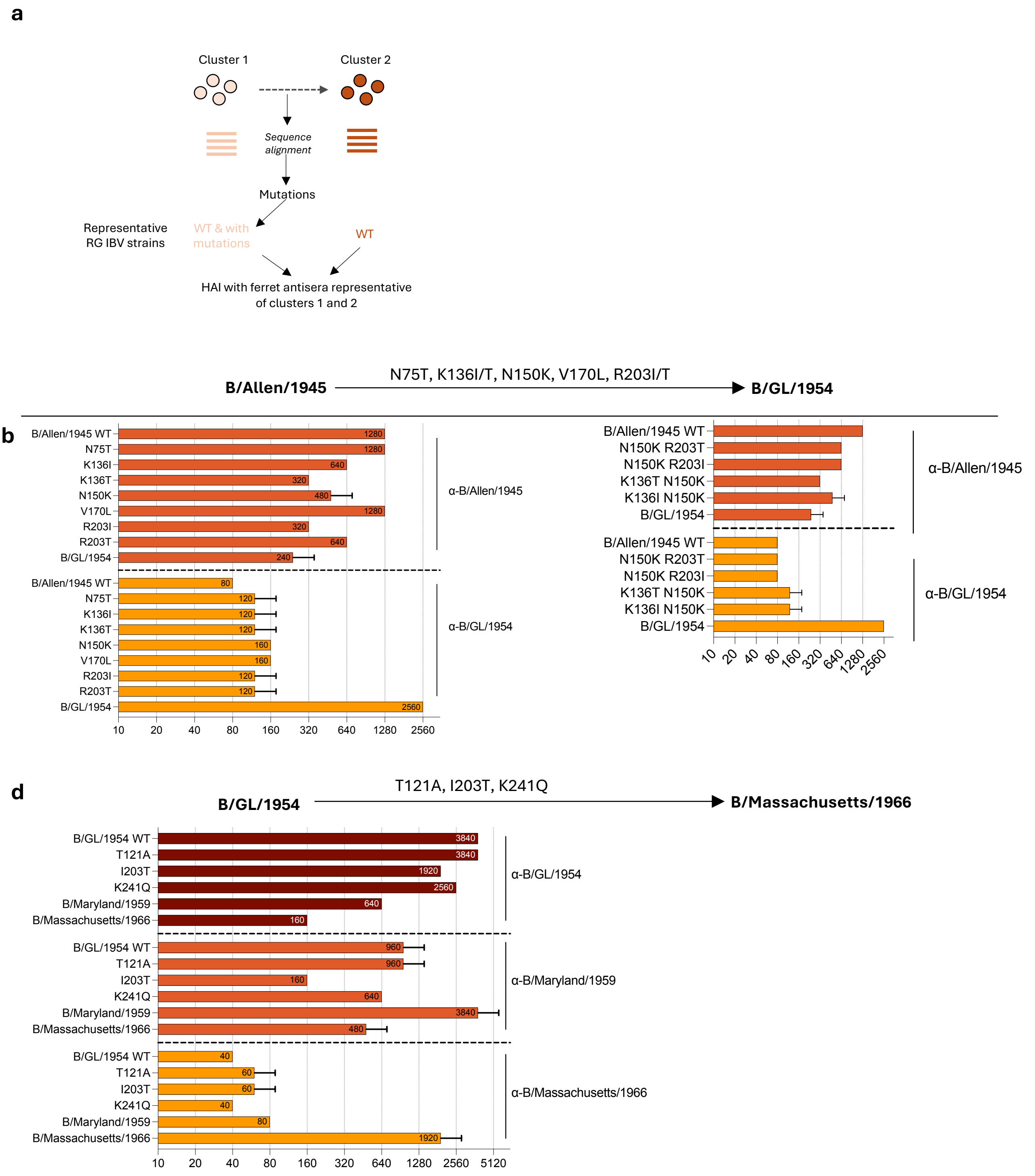


**Figure S3. Effect of amino acid substitutions on B/Allen/1945 and B/GL/1954 virus. (A)** Schematic summary of experimental approach. **(B)** Haemagglutination inhibition (HAI) activity of ferret antisera against viruses with indicated single or double mutations in HA on B/Allen/1945 background was tested against B/Allen/1945 and B/GL/1954 antisera. **(C)** B/GL/1954 mutations were tested against B/GL/1954, B/Maryland/1959, and B/Massachusetts/1966 antisera. Data represent mean from duplicates, performed in one experiment.

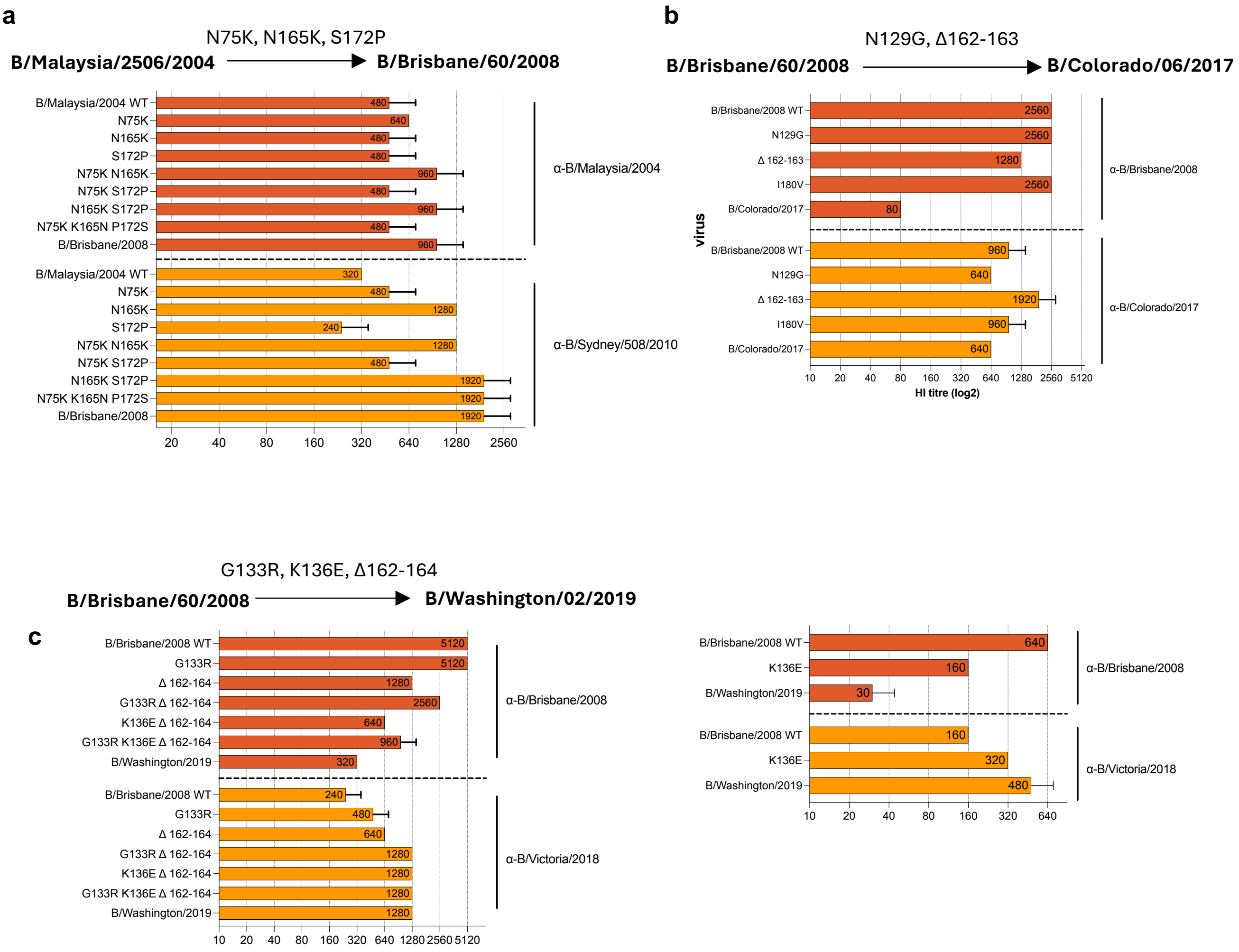


**Figure S4 Effect of amino acid substitutions on B/Victoria antigenic clusters.** Haemagglutination inhibition (HAI) activity of ferret antisera against viruses with mutations in HA on B/Malaysia/2506/2004 (a) and B/Brisbane/60/2008 (b-c) background were tested. Data represent mean from duplicates, performed in one experiment.

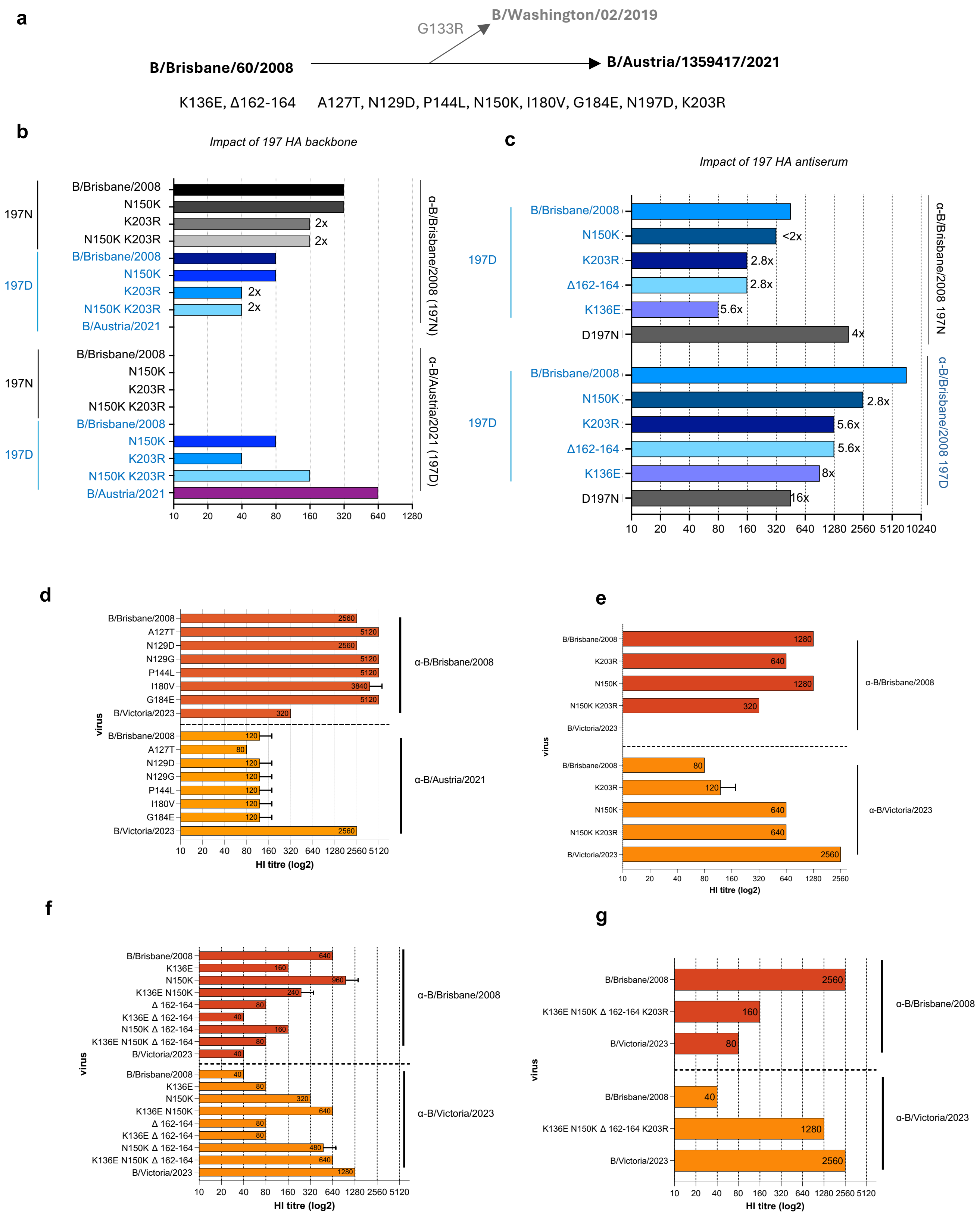


**Figure S5. Effect of amino acid substitutions on B/Brisbane HA. (a)** Mutation between B/Brisbane/60/2008 and B/Austria/1359417/2021 HA. **(b)** Impact of 197 HA backbone. The N150K and K203R mutations were introduced on the B/Brisbane/60/2008 197N and 197D HA backbones and the haemagglutination inhibition (HAI) activity of ferret antisera were tested. **(c)** Impact of 197 HA antiserum. The K136E, N150K and K203R mutations as well as the Δ162-164 deletion were introduced on the B/Brisbane/60/2008 HA and the haemagglutination inhibition (HAI) activities of ferret antisera raised against B/Brisbane/60/2008 197N or 197D HAs were tested. **(d-g)** Effect of mutations on the B/Brisbane/60/2008 HA background. Mutations were introduced on the B/Brisbane/60/2008 HA and the haemagglutination inhibition (HAI) activities of B/Brisbane/60/2008 and B/Austria/1359417/2021 or B/Victoria/16/2023 (B/Austria/1359417/2021-like) ferret antisera were tested. Data represent mean from duplicates, performed in one experiment.

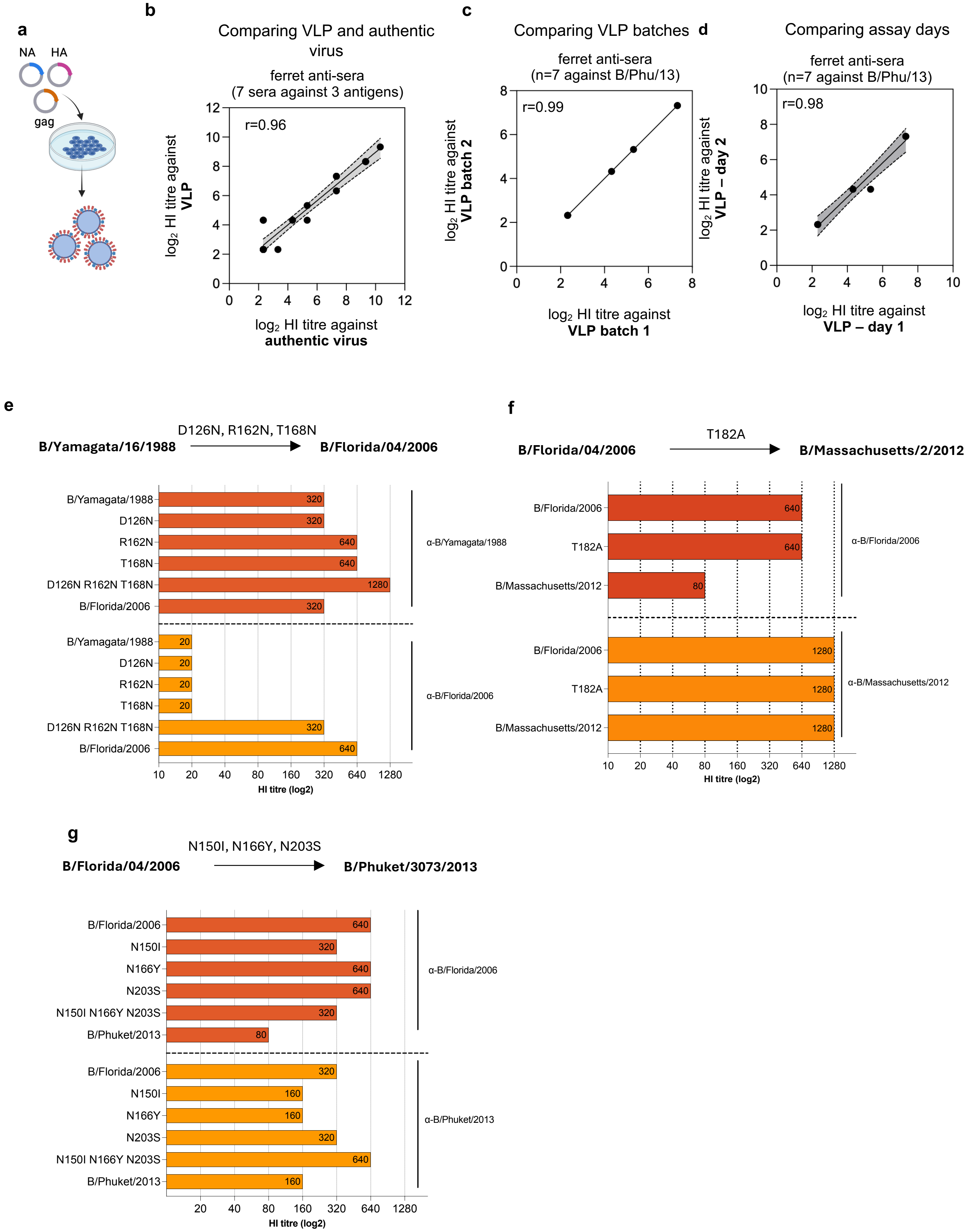


**Figure S6.** Validation of VLP-based HI assay and effect of amino acid substitutions in B/Yamagata. **(a)** Schematic of IBV VLP generation for HAI assays. Validation of VLP HAI assay by comparing HAI titers against authentic virus **(b)**, different batches of VLPs **(c)** and assays performed on different days **(d)**. Haemagglutination inhibition (HAI) activity of ferret antisera against VLPs with mutations in HA on B/Yamagata/16/1988 (e) and B/Florida/04/2006 (f-g) background were tested. Data represent mean from duplicates, performed in one experiment.

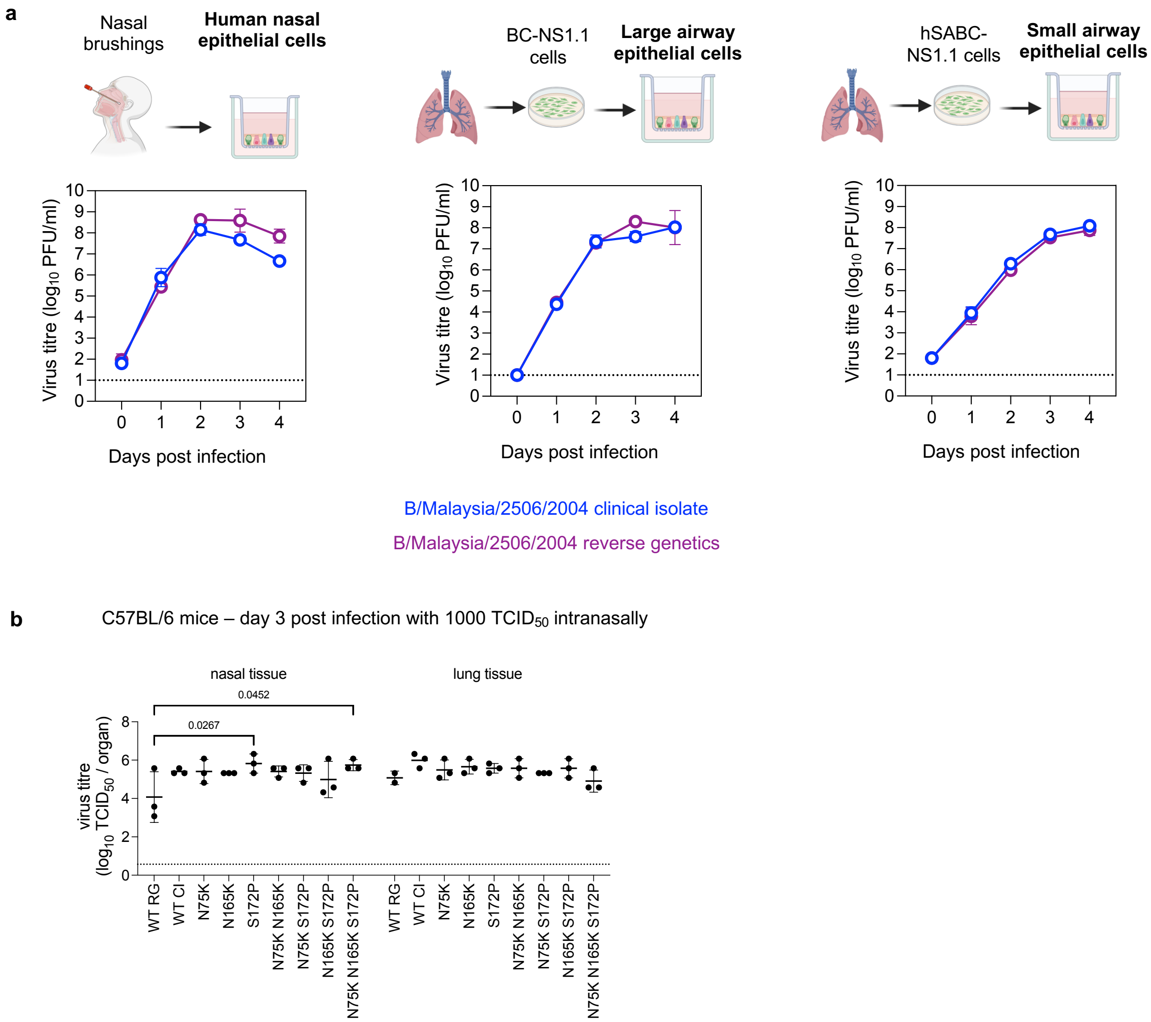


**Figure S7.** Replication of IBV in ALI differentiated human airway nasal epithelial cells (hNECs), large-airway epithelial cells (LAECs) or small-airway epithelial cells (SAECs) and mice. **(a)** Comparison of wildtype B/Malaysia/2506/2008 originating from a clinical isolate or rescued by reverse genetics. **(b)** Replication of B/Malaysia/2506/2004 WT and mutants in *in vivo*. Male C57BL/6 mice were intranasally infected with 1000 $TCID_{50}$ in 50 µL of virus. Tissues were harvested 3 days post infection, and viral titres were determined by $TCID_{50}$ assay. Dotted lines indicate the limit of detection. Mean and standard deviation (n=3 mice) from one experiment are shown. Statistical significance was determined by a 2-way ANOVA with Sidak's correction for multiple comparisons. RG refers to WT virus generated by reverse genetics and CI refers to WT virus that originated from a clinical isolate.
